# Deep neural networks trained for estimating albedo and illumination achieve lightness constancy differently than human observers

**DOI:** 10.1101/2025.07.10.664065

**Authors:** Alban Flachot, Jaykishan Patel, Thomas S. A. Wallis, Marcus A. Brubaker, David H. Brainard, Richard F. Murray

## Abstract

Lightness constancy, the ability to create perceptual representations that are strongly correlated with surface albedo despite variations in lighting and context, is a challenging computational problem. Indeed, it has proven difficult to develop image-computable models of how human vision achieves a substantial degree of lightness constancy in complex scenes. Recently, convolutional neural networks (CNNs) have been developed that are proficient at estimating albedo, but little is known about how they achieve this, or whether they are good models of human vision. We examined this question by training a CNN to estimate albedo and illumination in a computer-rendered virtual world, and evaluating both the CNN and human observers in a lightness matching task. In several conditions, we eliminated cues potentially supporting lightness constancy: local contrast, shading, shadows, and all contextual cues. We found that the network achieved a high degree of lightness constancy, outperforming three classic models, and substantially outperforming human observers as well. However, we also found that eliminating cues affected the CNN and humans very differently. Humans had much worse constancy when local contrast cues were made uninformative, but were minimally affected by elimination of shading or shadows. The CNN was unaffected by local contrast, but relied on shading and shadows. These results suggest that the CNN followed an effective strategy of integrating global image cues, whereas humans used a more local strategy. In a follow-up experiment, we found that the CNN could learn to exploit noise artifacts that were correlated with illuminance in ray-traced scenes, whereas humans did not. We conclude that CNNs can learn an effective, global strategy of estimating lightness, which is closer to an optimal strategy for the ensemble of scenes we studied than the computation used by human vision.

## 1 Introduction

Lightness constancy is the ability to create perceptual representations that are strongly correlated with the achromatic surface reflectance of objects, across a wide range of lighting conditions and contexts. Like color constancy, this feature of our visual system helps us to build a stable and robust representation of our environment (Anderson, 2011), which is essential for high level tasks such as object recognition (Murray, 2021; Hurlbert, 2007; Foster, 2011). However, despite a long tradition of empirical and theoretical studies, it remains unclear what strategy the visual system uses to perform this feat, and for a simple reason: lightness perception is complex and depends on many different features, from low to high-level, and from local to global. Some studies have shown the importance of local contrast in human lightness constancy (Gilchrist, 2006; Wallach, 1948, 1963; Adelson, 2000; Arend et al., 1991; Arend & Spehar, 1993; Foster, 2011; Kraft & Brainard, 1999; Wedge-Roberts et al., 2020). Others have shown that global contextual features can greatly influence human lightness and color judgment, including scene complexity (Kraft et al., 2002), object shading and shadows (Szafir et al., 2015), luminance and chromaticity range (Adelson, 1993; Webster & Mollon, 1995), and specular highlights (Wedge-Roberts et al., 2020). Additional studies have shown the relevance of high-level features such as memory and familiarity, particularly in the natural world (Tsuda et al., 2020; Granzier & Gegenfurtner, 2012). A consequence is that despite many attempts (Land & McCann, 1971; Funt et al., 2000; Dakin & Bex, 2003; Murray, 2021), an accurate and image computable model of this phenomenon has yet to be successfully developed.

Recent studies have shown that deep-learning neural networks (DNNs) are a promising approach to performing many visual tasks (Krizhevsky et al., 2012; Yao et al., 2020; Storrs et al., 2021). In particular, DNNs have made tremendous progress towards solving illumination estimation for complex and naturalistic images (Afifi et al., 2022; Ulucan et al., 2024). Furthermore, deep learning models for intrinsic image decomposition have been successful at extracting the physical surface albedo of objects in complex scenes (Heidari-Gorji & Gegenfurtner, 2023; Z. Li & Snavely, 2018b; Z. Li et al., 2020; Wang et al., 2021; Fan et al., 2018; Yu & Smith, 2019). DNNs trained for color classification and color constancy have also been shown to develop supra-human levels of constancy and human-like representations of color (Flachot et al., 2022). And recently, Murray et al. (2023) showed how these models can be tested for lightness constancy on the same images as human observers, allowing for a direct and behaviorally relevant comparison. DNNs are therefore interesting candidates for producing a useful, image computable model of human lightness constancy.

In this study, we aim to extend our understanding of human lightness constancy via a series of experiments, while exploring the potential of a recent and promising family of deep learning algorithms as models of human lightness constancy. We aim to answer three main questions: (1) What features do human observers rely on for lightness constancy in complex scenes? (2) Can DNNs, and in particular those trained for intrinsic image decomposition, achieve human levels of lightness constancy on the same images? (3) And perhaps most importantly, do these models use the same features as human observers?

To answer these questions, we devised a lightness matching experiment and rendered images of three-dimensional scenes to test both human participants and DNNs trained for intrinsic image decomposition. In several different conditions, we manipulated cues in the rendered scenes that previous work has found to be relevant to lightness and color constancy for human observers (Gilchrist, 2006; Szafir et al., 2015; Kraft & Brainard, 1999). This allowed us to identify what features human observers and DNNs rely on for lightness constancy.

We found that the models were able to achieve a high degree of lightness constancy in complex scenes. However, the features the models most relied on differed substantially from those used by human observers. Furthermore, the features used by DNNs varied depending on the rendering engine used to generate training and test images, but did not vary in this way for human observers.

## 2 Experiment with human observers

### 2.1 Methods

#### 2.1.1 Observers

Nine observers participated in the experiment. All were naive except one, who was the first author. Five were male, four were female, and ages ranged from 25 to 34 years. All reported normal or corrected-to-normal vision, and gave written informed consent. All procedures were approved by the Office of Research Ethics at York University.

#### 2.1.2 Stimuli

We rendered 256 × 256 pixel images with three color channels (RGB) using the EEVEE engine in Blender 2.92 (Fig. 1(A)) (Blender 2.92 Documentation Team, 2021). When describing rendered stimuli, we use the term ‘RGB albedo’ to mean the triplet of simulated reflectances assigned to a surface in the three color channels, in the scene description provided to the renderer, and ‘achromatic albedo’ or simply ‘albedo’ to mean the simulated reflectance in a single channel. The rendered scenes had two walls and a floor, with 14 objects of different shapes, RGB albedos, sizes, and positions, randomly sampled from the distribution we used for training deep learning networks (see Section 3.1). Each image showed the same arrangement of geometric objects. The RGB albedo assigned to each object was the same in all images, except for a small number of test objects, where it varied from image to image as described below. In order to speed rendering and introduce some variability into the stimuli, we rendered RGB images with different randomly chosen albedos in the three color channels, and used each channel of the rendered image as a separate achromatic stimulus in the experiment. As a result, there were three subsets of stimulus images that showed objects with different achromatic albedos (Fig. 1(A)). We refer to each single-channel image as a ‘luminance’ image, since it was used to generate the on-screen luminance. (Thus the luminance image was not a weighted sum of the color channels, as is sometimes the case.) For each RGB image, Blender also rendered an image that gave the RGB albedo at each pixel. *Illuminance* is incident luminous flux per unit area (e.g., lumens/m^2^ in SI units), and for a Lambertian surface it is proportional to the ratio of luminance and reflectance (McCluney, 2014). For each stimulus we calculated an ‘illuminance image’ that indicated the amount of incident light at each location, by taking the ratio of the luminance and achromatic albedo images at each pixel.

**Figure 1:**
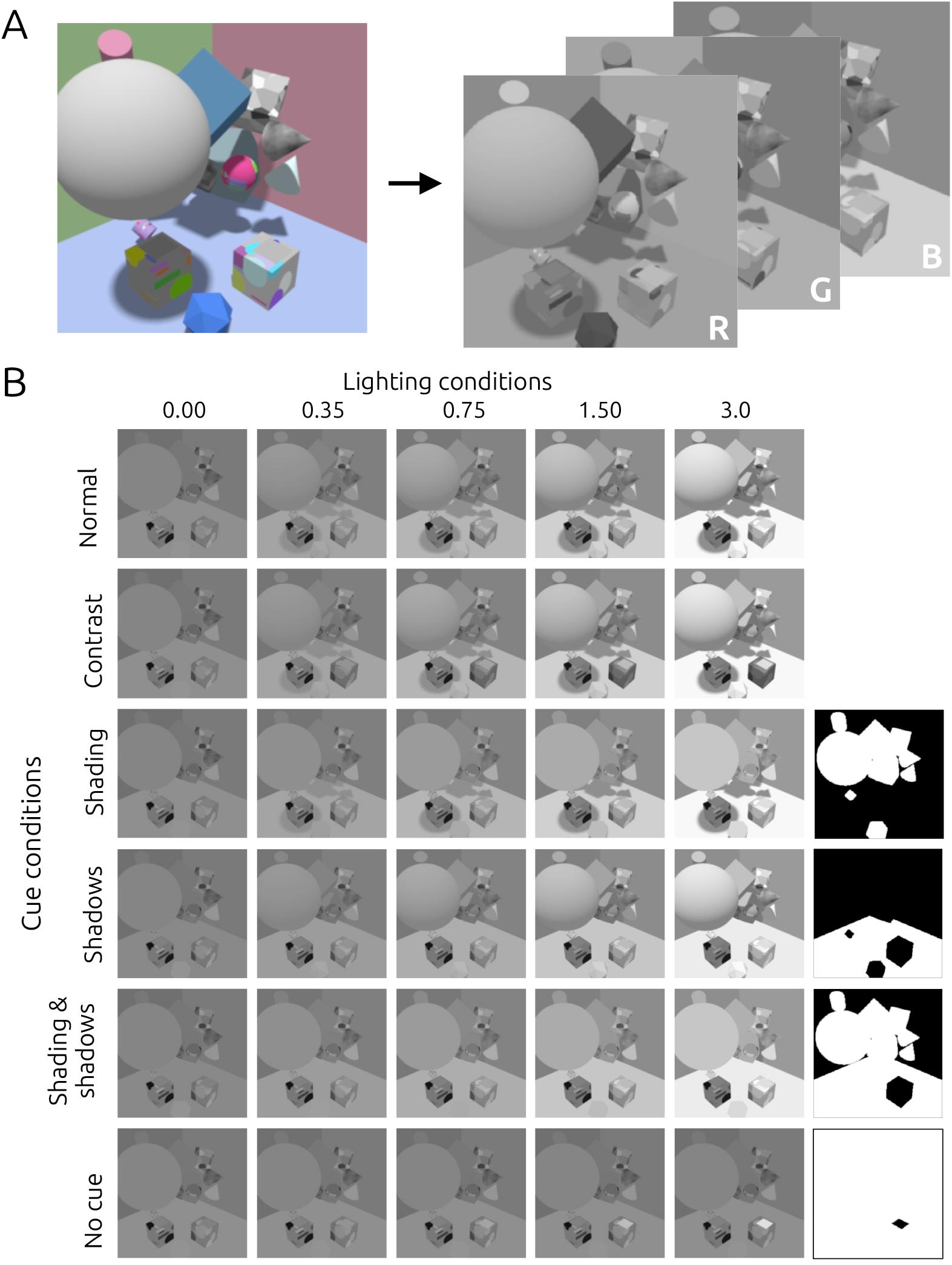
Stimulus images in the lightness matching experiment. (A) A typical image, rendered in RGB and converted into three achromatic images, one from each color channel. (B) Lighting and cue manipulations. Each column shows stimuli in a single lighting condition, with the indicated value of point light intensity. Each row shows stimuli in a different cue condition, where we manipulated contrast, shadow, and shading cues. The black-and-white images to the right of the bottom four rows are masks that we used to create the stimuli in these conditions, as described in the main text.

Each scene had two achromatic light sources: an ambient light, with intensity fixed at 0.35 in Blender’s intensity units (IU), and a point light located above and outside the visible scene, with intensity set to 0.00, 0.35, 0.75, 1.50 or 3.00 IU, as shown in Fig. 1(B). The cube under the large sphere was the *reference cube*, and the one to the right was the *match cube*. On top of the reference cube was the *reference patch*, and on top of the match cube was the *match patch*. The reference patch albedo was 0.20, 0.40 or 0.60. The match patch albedo ranged from 0.00 to 1.00, in increments of 0.025. We rendered stimuli with all combinations of these scene parameters: five point light intensities, three reference patch albedos, and 41 match patch albedos. To vary the availability of cues that could potentially support lightness constancy, we created six variations of these stimuli (Fig. 1(B). (1) In the Normal condition, the stimuli were rendered as described above. (2) In the Contrast condition, the albedo of the match cube was chosen such that the average luminance of the top face of the match cube (surrounding the match patch) was the same in every image, and equal to the average luminance of the top face of the reference cube (surrounding the reference patch). As a result, matches based on local contrast would equate the luminance of the reference and match patches, not their albedos. This manipulation silenced local contrast as a cue to albedo.

In the remaining four conditions, the stimuli were created by taking images from the Normal condition, and replacing a subset of pixels by corresponding pixels from Normal images where only ambient light illuminated the scene (Fig. 1(B), top left image, labelled Normal and 0.00). The white pixels in the black-and-white images on the right in Fig. 1(B) indicate which pixels were replaced in each condition. The values of the inserted pixels were scaled so that their mean was the same as that of the pixels they replaced. (3) In the Shading condition, pixels depicting the 14 random geometric objects were replaced by pixels from ambient-illuminated images. This eliminated shading and shadows on the 14 objects. Cast shadows on the walls and floor were preserved. (4) In the Shadows condition, all pixels depicting the floor were replaced by corresponding pixels from ambient-illuminated images. This eliminated cast shadows on the floor. (5) In the the Shading & Shadows condition, all pixels depicting the 14 geometric objects and the floor were replaced by pixels from ambient-illuminated images. Only pixels depicting the reference and match cubes and the walls were left unchanged. Finally, (6) in the No-cue condition, all pixels except those depicting the reference and match patches were replaced by corresponding pixels from ambient-illuminated images. This was a limiting condition, where there were no cues to the simulated lighting, and in this condition we expected observers to simply match luminance instead of albedo, i.e., to show no lightness constancy. The average luminance around the reference and match patches was the same in the Contrast and No-cue conditions.

The stimuli were back-projected onto a 100 cm wide × 70 cm high screen with a ProPixx 1440 Hz projector in a dark room. Observers sat 120 cm from the screen, and head position was stabilized with a chinrest. The projected stimulus image was 25 cm square, and subtended 11.9 degrees of visual angle, horizontally and vertically. The image was surrounded by a grey background projected onto the screen. We made luminance characterization measurements from the display using an Xrite i1Display Pro colorimeter, fitted a gamma-correction model, and transformed the stimuli using custom software so that displayed luminance was proportional to rendered luminance (Brainard et al., 2002). We scaled the rendered stimuli before display so that luminance ranged up to 125.4 cd/m^2^. We created the experiment using Python 3.8 and PsychoPy 2024.1.4 (Pierce et al., 2019). Python code for generating stimuli, running the experiment, and analyzing the results can be found in our GitHub repository ^1^.

#### Procedure

On each trial, the observer viewed an achromatic stimulus randomly chosen from one of the six cue conditions (Normal, Contrast, Shading, Shadows, Shading & Shadows, and No-cue). Images from the three rendered color channels were randomly interleaved. The simulated point light intensity was randomly set to 0.00, 0.35, 0.75, 1.50 or 3.00 IU. The reference patch albedo was randomly set to 0.20, 0.40, or 0.60. The match patch albedo was randomly set to an initial value between 0.00 and 1.00. The observer was instructed to match the appearance of the match patch to that of the reference patch. The observer used a mouse scroll wheel to adjust the albedo of the match patch, in steps of 0.025 albedo units, until it appeared to have the same albedo as the reference patch. For each albedo of the match patch, the software loaded and displayed a new rendered image. There was no limit on response time, although the observer was told they would be evaluated based on both accuracy and speed, so they were incentivized to make quick, visual match settings instead of cognitively inferred settings. When the observer was satisfied with the match, they clicked the mouse, and after a brief pause (0.3 s) the next trial began.

There were six conditions, five point light intensities, three reference albedos, and three color-channel to grayscale conversions, for a total of 6 × 5 × 3 × 3 = 270 trials. These trials were interleaved with additional trials showing similar stimuli rendered with the Cycles engine in Blender 2.92, for a comparison we describe below; see Section 4. Despite the inter-trial interval of 0.3 s, it occasionally happened that the mouse button press that ended one trial had not been released when the next trial began, in which case the initial, randomly chosen match albedo was saved as a match, with a response time near zero. These trials amounted to 1.5% of the total, and were not included in the analyses. On average, valid trials had a response time of 7.1 s, and the whole experiment took 42 minutes.

#### Analysis

We quantified lightness constancy using the Thouless ratio (Thouless, 1931), defined as follows. Let *r*_1_ be the albedo of the reference patch in a matching task, and let *r* be the albedo that the observer chooses at the match patch. Let *i*_1_ and *i* be the illuminances at the reference and match patches, respectively. An observer who simply matches luminance (i.e., has no lightness constancy) chooses the match albedo *r*_0_ where *r*_0_*i* = *r*_1_*i*_1_, which implies *r*_0_ = *r*_1_*i*_1_/*i*. The Thouless ratio *τ* is defined as

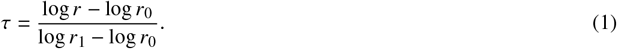

A perfectly lightness-constant observer (*r* = *r*_1_) has a Thouless ratio *τ* = 1, and a luminance-matching observer (*r* = *r*_0_) has *τ* = 0. The logarithms in this definition reflect the fact that perceived lightness is a monotonic, compressive function of physical albedo. The Thouless ratio is similar to the color constancy index that is often used to quantify constancy in a multi-dimensional color space (Arend et al., 1991; Brainard, 1998).

Equation (1) can be rewritten as

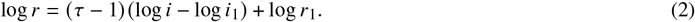

This shows that for an observer with a fixed Thouless ratio *τ*, log match albedo (log *r*) is an affine function of log match illuminance (log *i*), with slope *τ* − 1. When the observer’s albedo match is invariant across different illumination intensities (perfect constancy, *τ* = 1) the slope is 0. When the observer matches luminance (no constancy, *τ* = 0), the slope is −1. We estimated each observer’s Thouless ratio in each condition from the slope of a least-squares linear regression line fitted to the logarithm of the match albedo settings as a function of the logarithm of illuminance at the match patch.

### 2.2 Results

Fig. 2(A) shows results for a typical observer in all conditions. Each panel plots match albedo against match illuminance (represented as a multiple of the illuminance at the reference patch), both on a log scale. Each color shows results for a single reference albedo. Log match albedo was an approximately linear function of log illuminance: across all observers, conditions, and reference albedos, the matches were well fit by linear regression, with an average coefficient of determination R^2^ = 0.92. The bracketed values on each line are the Thouless ratios estimated from the line’s slope as in equation (2).

**Figure 2:**
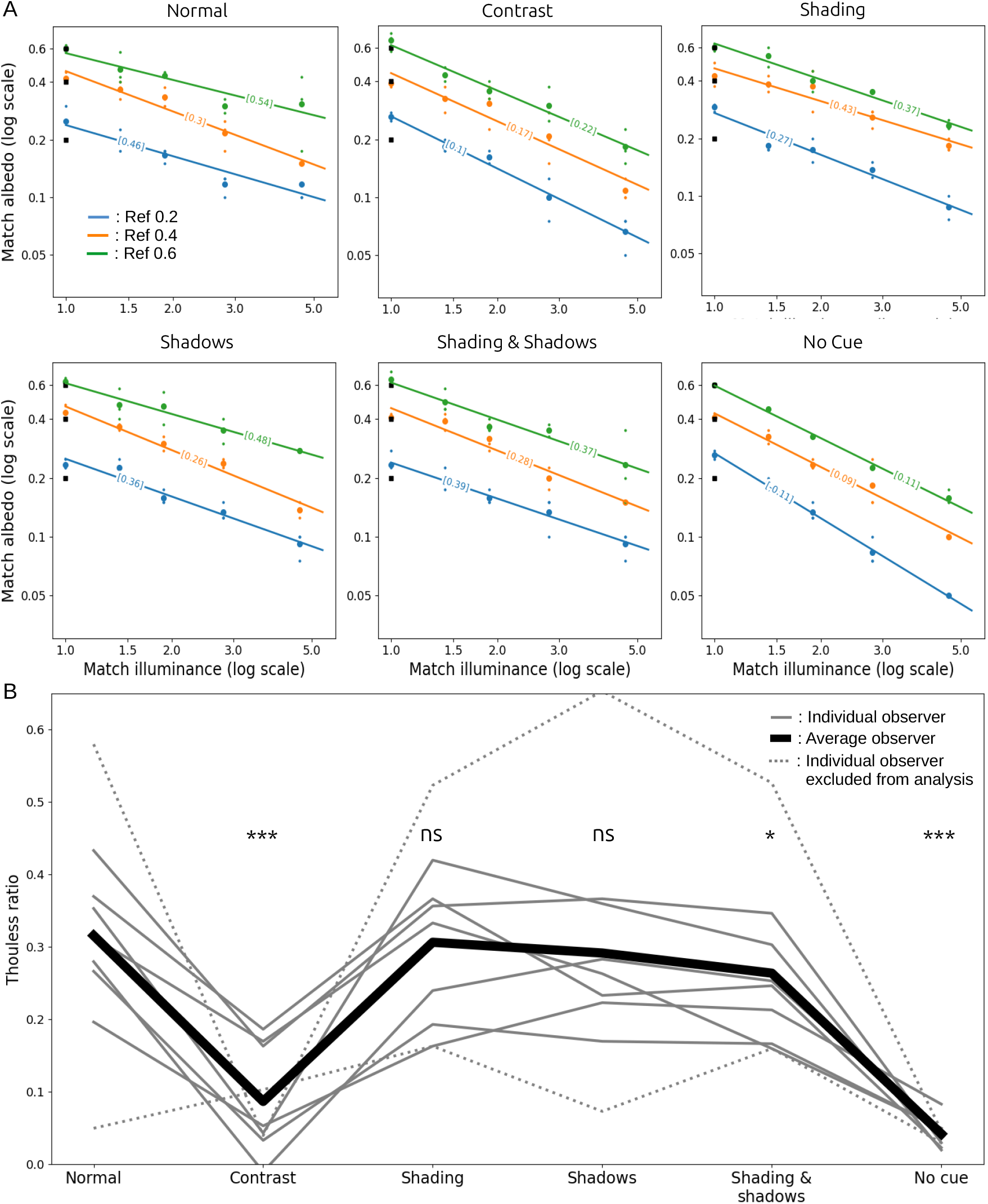
(A) Results for a typical observer in all conditions. Log match albedo declined as an approximately linear function of log match illuminance. Bracketed values show Thouless ratios estimated from the fitted lines. (B) Thouless ratios for individual observers (grey lines) and the mean across observers (black line). Significance levels are the results of paired *t*-test comparisons to the Normal condition. (∗ for *p* < 0.05, ∗∗ for *p* < 0.01 and ∗. ∗ ∗ for *p* < 0.001)

Constancy was approximately the same for the three reference albedos. A one-way repeated measures ANOVA testing for an effect of reference albedo on Thouless ratio did not reach significance (*F* (2, 12) = 0.53, *p* = 0.69). However, matches were slightly different for images from the three color channels. A one-way repeated-measures ANOVA of the effect color channel on match reflectance was significant (*F* (2, 12) = 69.8, *p* < 10^−6^). Nevertheless, the differences were small: the average matches for the R, G and B channel stimuli were 0.25, 0.26, and 0.28, respectively. Small differences in lightness matches between these groups of stimuli are not surprising, as they had different albedos assigned to walls and objects.

Fig. 2(B) shows Thouless ratios for all observers in all conditions. Light grey lines show results for individual observers, averaged over reference albedos, and the heavy black line shows the average over observers and reference albedos. The two dotted lines show results for two observers we excluded from the analysis. The lower dotted line indicates poor constancy in all conditions, and so the observer may not have understood the task. The upper dotted line was from the first author, a highly practised and non-naive observer who had unusually good constancy in some conditions (but nevertheless a qualitatively similar profile across conditions, compared to the other observers).

Overall, Thouless ratios were relatively low, with an average of 0.32 in the Normal condition. It is not unusual to find weak constancy when stimuli are not rich, naturalistic scenes (Katz, 1935). Here the stimuli were artificial Lambertian scenes, with simple lighting conditions, shown on a 2D display, and these factors may have contributed to relatively weak constancy.

The pattern of Thouless ratios across conditions was quite consistent across observers: highest constancy in the Normal condition; little change in the Shading, Shadows, and Shading & Shadows conditions, suggesting that observers were largely indifferent to object shading and shadows (after the experiment, several observers reported being unaware that shadows were sometimes absent); lowest constancy in the No-cue condition, as expected, with an average Thouless ratio of 0.04; and low constancy in the Contrast condition, with an average Thouless ratio of 0.09. These observations strongly suggest that, for our stimuli, the perceived color of the match patch is mostly influenced by local contrast. The asterisks in Fig. 2(B) show the results of paired *t*-tests that compared each condition to the Normal condition, which support these qualitative observations.

Although there were individual differences, most observers exhibited a similar behavioral pattern, namely low sensitivity to shading and shadows, and high sensitivity to local contrast. In the following section, we test whether the same results can be found with a deep neural network trained to estimate the albedo of objects independently of lighting conditions and context.

## 3 Deep learning models

We trained a deep learning network for intrinsic image decomposition, and evaluated it for lightness constancy by simulating its performance on same task we studied with human observers.

### 3.1 Training images

We rendered 121K training images and 7K validation images using the EEVEE engine in Blender 2.92 (Blender 2.92 Documentation Team, 2021). Each scene included 17 randomly chosen geometric shapes (spheres, rectangular prisms, cylinders, tori, and icosahedra), with randomized locations, orientations, and sizes (Figure 3). Each object was rendered with a surface pattern that was randomly chosen to be (a) a solid colour, with an albedo uniformly sampled between 0.05 and 0.9, which is approximately the range of albedos found in natural scenes (Gilchrist, 2006), (b) a Voronoi pattern, (c) a low-pass noise pattern or (c) a random composite of filled-in rectangles and ellipses, again with albedo uniformly sampled between 0.05 and 0.9. The scenes were illuminated by an ambient light source with a random intensity and a directional light source with random direction and intensity. Both light sources had randomized colours. The virtual camera was placed at a randomly chosen location and was directed at the centre of the scene. We also rendered ground-truth albedo images of each scene, and calculated illuminance images as the pixelwise ratio of luminance and albedo images. Images were saved in EXR format with 32-bit floating point precision. The Supporting Information includes Python code that generates these images and provides further details about randomized parameters.

**Figure 3:**
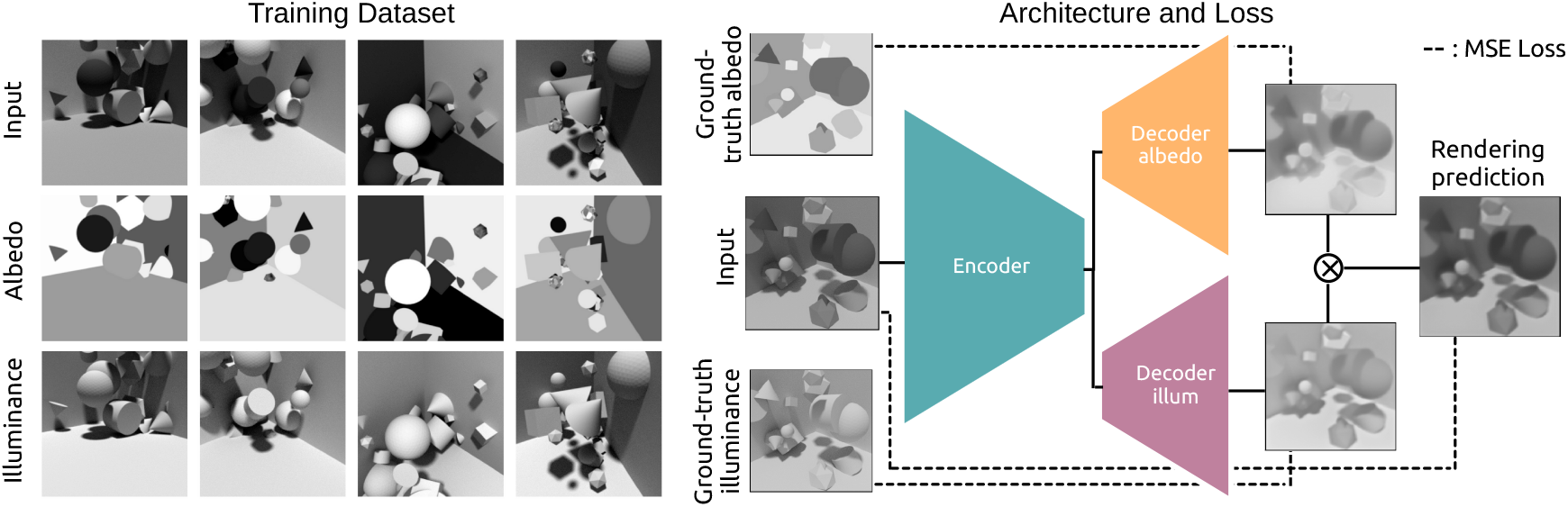
Training procedure for the convolutional neural network. Left: Examples of training images. Right: Architecture and loss.

We rendered the images in colour, but used the three colour channels as separate achromatic training images. We also augmented the training data using vertical and horizontal flips. These augmentations produced a training set with 121K × 3 channels × 4 flips = 1.45M achromatic images for each of luminance, albedo, and illuminance.

### 3.2 Model, loss, and training

The network had a modified U-Net architecture with one encoder, and two separate decoders for albedo and illuminance (Fig. 3(B)) (Z. Li et al., 2020). The encoder had 2240 convolutional filters split over six convolutional layers. Both decoders had 1408 convolutional filters, also split over six convolutional layers. Each layer of the encoder sent its output to the next encoder layer in a feedforward manner, and also to the decoders via direct skip connections.

We trained the model to map training images to their corresponding ground-truth albedo and illuminance images. The cost function for each estimate was

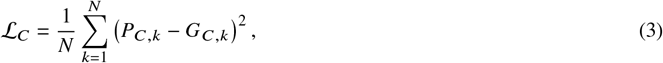

where *P* is the model’s prediction, *G* is the ground-truth image, *C* stands for either *albedo* or *illuminance*, and *N* is the number of pixels in the image.

Because the scenes depicted Lambertian surfaces, each input image *I* was the product of its two intrinsic components, albedo and illuminance. As a regulatory term, we imposed a reconstruction cost by penalizing any discrepancy between the input image and the pointwise product of the model’s albedo and illuminance predictions. We defined the reconstruction loss as

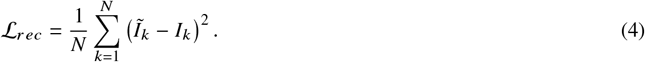

where 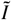 is the pixelwise product of the model’s estimates of albedo and illuminance, *I* is the input image, and the sum is over image pixels.

The total loss ℒ_*R*_ was a combination of three terms:

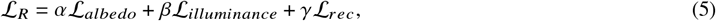

where *α, β*, and *γ* are scalars. Because we evalauted models on their ability to estimate albedo, we favored learning albedo estimation over the other criteria by setting *α* = 4 and *β* = *γ* = 1. The illuminance and reconstruction losses served as regulatory cost functions.

Training lasted for 60 epochs, and in each epoch the network was trained on 125K images × 3 channels = 375K images. Each image was flipped horizontally and vertically with probability 0.5 for each direction. We used the Adam optimizer with a learning rate that started at 5 × 10^−5^ and decayed by a factor of two every ten epochs. The loss flattened out around the 50th epoch. The Supporting Information provides code that implements the network and training routines.

Because the training procedure included several random steps, such as the initialization of the weights and the order in which images were shown, different training runs resulted in different learned weights. We performed three training runs, resulting in three different training instances.

### 3.3 Validation

After training, we evaluated the network’s ability to estimate albedo on the validation set, which consisted of images that were generated by the same random procedure that created the training images, but that were not used during training. We found that the albedo output of the network largely removed shading and shadows from input images (Fig. 4). Furthermore, the network produced generally good pixelwise albedo estimates, with a median absolute error of 0.028, measured in albedo units that range from 0 to 1 (Fig. 5).

**Figure 4:**
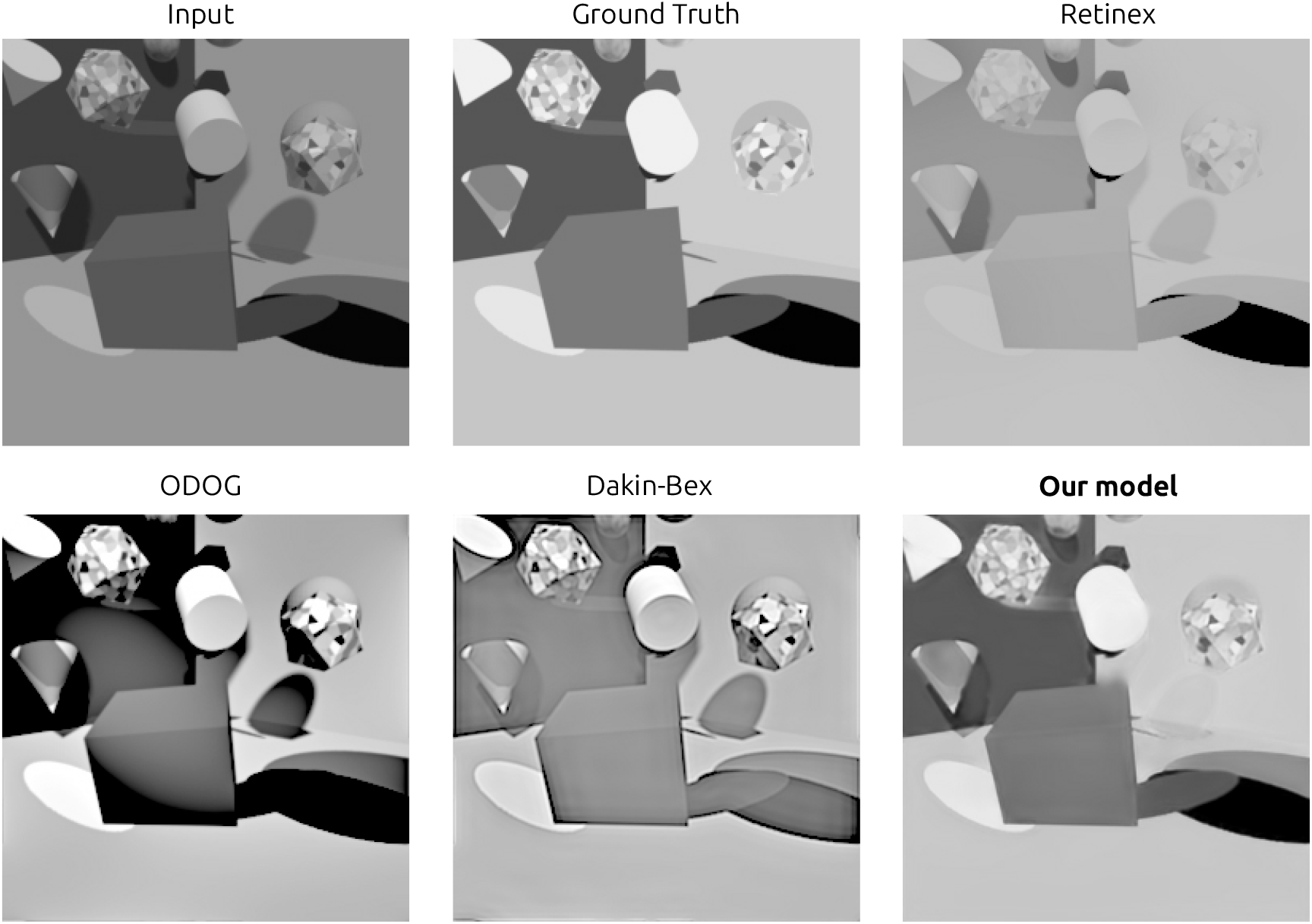
Example of an image from the validation set, its ground-truth albedo, and the predictions from the models tested. The deep neural network’s prediction were the closest to the ground truth, and it was the only model able to largely eliminate shading and shadows.

**Figure 5:**
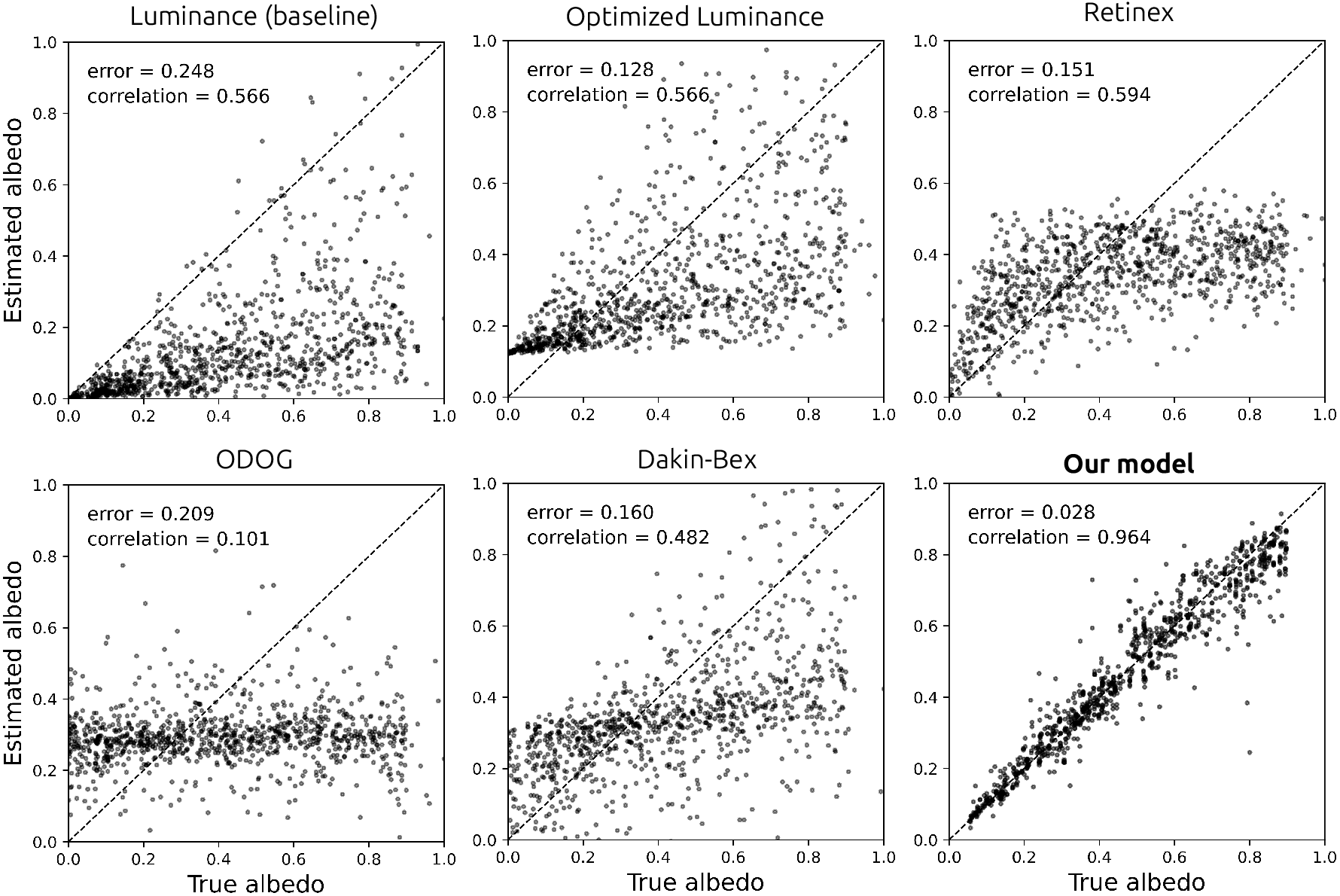
Scatter plots of models’ albedo estimates against the ground-truth albedo. The DNN model had the best performance of the models tested. The reported errors are the median absolute prediction error across pixels.

For comparison, we also evaluated a number of alternative models. There are few image-computable lightness models in the psychophysics literature, so we evaluated models of both lightness (perceived albedo) and brightness (perceived luminance).

We evaluated ODOG, which is a brightness model (Blakeslee & McCourt, 1999), retinex, which is a lightness model (Land & McCann, 1971; Funt et al., 2000), and the Dakin-Bex normalization model, which has been described in both ways (Dakin & Bex, 2003). These models do not give outputs in albedo units, so we optimized their performance by finding the affine transformation that minimized each model’s median absolute error when predicting albedo in the training set, as follows. We applied each model to the 128K images in the training set. We found the affine transformation that gave each model the lowest median absolute error when predicting albedo at 10^3^ pixels randomly sampled from each training image, for a total of 1.28 × 10^8^ pixels. We then used these optimal affine transformations, estimated from the training set, to map model outputs to predicted albedos on the validation set. As baselines, we also tested two null models whose outputs were in one case simply the luminance input, and in the other case the luminance output modified by an optimal affine transform as with the other models.

Unlike the deep learning network, the outputs of these alternative models generally did not remove shading and shadows (Fig. 4). Furthermore, even after an optimal affine transform of their outputs, they had substantially higher prediction error than the deep learning network (Fig. 5). In fact, the three lightness/brightness models (ODOG, Dakin-Bex, and retinex) gave albedo estimates whose median absolute errors were even higher than that of the optimized luminance model, whose output was simply an affine transformation of the luminance input. Patel et al. (submitted) found similar results for a trained network and classical models, using a similar but not identical network and training set.

The validation results show that training was successful, in that the network was able to decompose rich 3D scenes into their albedo and illuminance intrinsic image components, at least when the scene was typical of the training distribution. We also found that the network performed better than classical models of lightness and brightness. As such, the network is an interesting candidate for a model of human lightness constancy. In the following section we evaluate the network in the same matching task we used with human observers, for a more direct comparison.

### 3.4 Lightness constancy in a matching task

To further evaluate the network’s degree of lightness constancy, we ran it in a simulation of the matching task performed by human observers in the behavioural experiment reported above. In each condition of the simulated experiment, and for each lighting configuration and reference albedo, we found the network’s match setting as follows. We applied the network to the same stimuli viewed by human observers, with a fixed reference patch albedo and with a range of match patch albedos. We took the network’s match setting to be the match patch albedo for which the network’s mean output was the same at the reference patch and the match patch. For the finite set of test stimuli, the model’s output was never exactly the same at the reference and match patches, so we found the match setting by interpolation or extrapolation. We then calculated a Thouless ratio for the network, just as we did for human observers, from a fit of equation (2) to the logarithm of the network’s match albedo setting as a function of log illuminance at the match patch.

In all stimulus conditions, the networks’ log match albedo was an approximately linear function of log illuminance (Fig. 6A). This is itself a noteworthy finding, and an important similarity to human observers that did not have to be the case for such a complex and highly nonlinear network. This linearity also means that the Thouless ratio is a meaningful metric for describing the network’s performance in the matching task. The Thouless ratio for each condition is shown in brackets on each fitted line.

**Figure 6:**
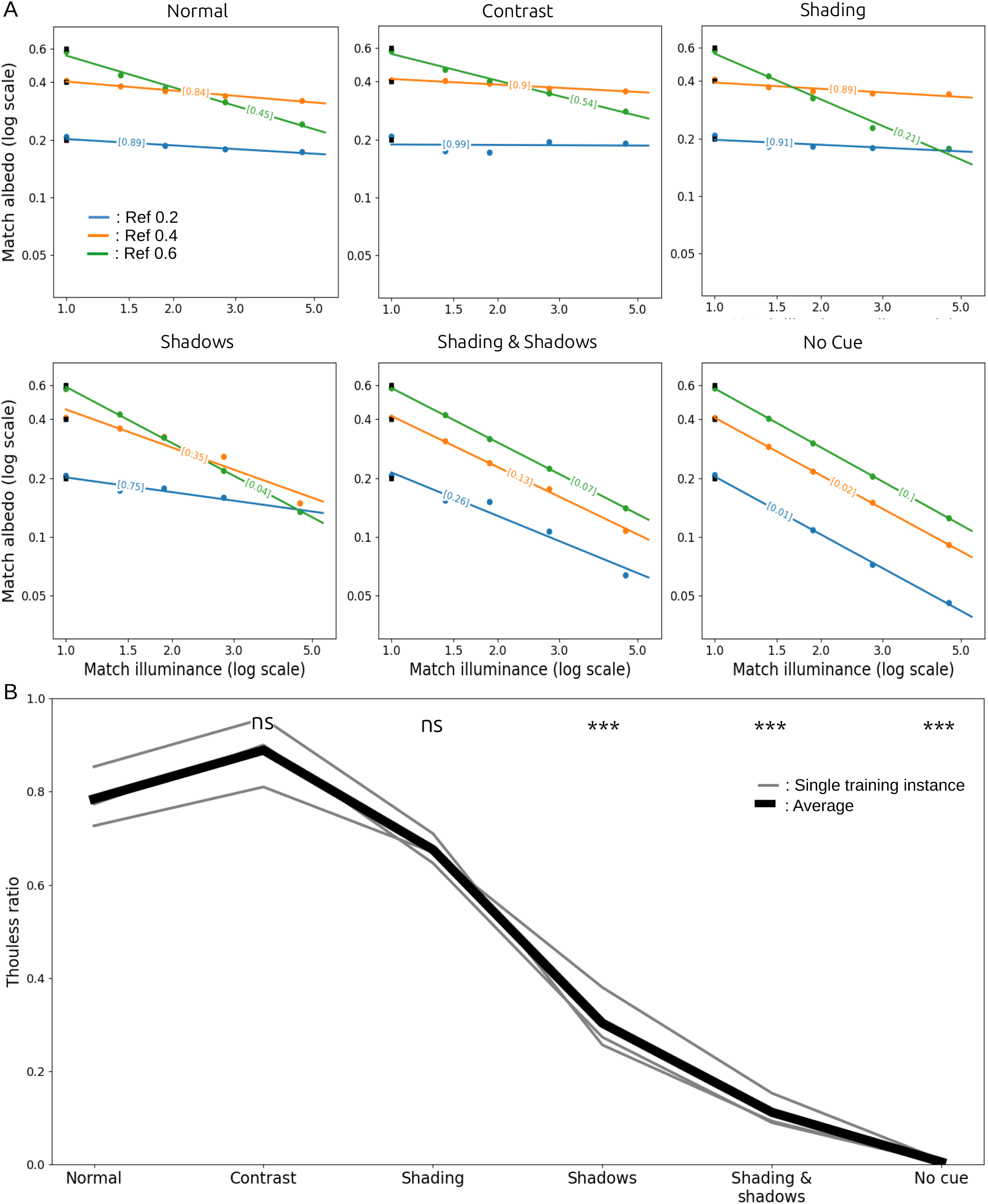
(A) Results for one training instance of the network in all conditions. (B) Thouless ratios for single training runs (grey lines, N = 3) and the average across training runs (black line). Results from different training runs were similar, and tend to overlap in the figure. Significance levels are the results of paired *t*-test comparisons to the Normal condition.

Fig. 6(A) also shows that this training instance’s match settings were largely constant as a function of illuminance in the Normal and Contrast conditions, and that they varied with illuminance in the remaining conditions. This differs from what we found for human observers. Additionally, the Thouless ratios varied markedly with the reference albedo (a one-way repeated measures ANOVA revealed an effect of albedo on match reflectance; *F* (2, 12) = 102.3, *p* = 0.0003), with much weaker constancy when the reference albedo was 0.6. A partial explanation of this difference can be found in the network’s illumination predictions (not shown here). When the reference albedo was 0.6, the model assigned an unusually high illuminance to the reference cube. What led the network to make this mistake, however, is unclear.

Fig. 6(B) shows the mean Thouless ratio for each condition, averaged across reference albedos for each training instance (grey lines) and averaged across training instances and reference albedos (bold black line). As was the case for human observers, the network’s constancy was affected by the contextual cue manipulations: a one-way repeated measures ANOVA revealed a significant effect of cue condition on Thouless ratio (*F* (5, 10) = 138.7, *p* ≪ 0.01). However, the similarity ends there, as some differences clearly emerge. First, in the Normal condition, the Thouless ratio was much higher for the network (mean 0.79) than for human observers (0.38), indicating that the network had markedly better lightness constancy than humans in the realistic baseline condition. (Note that the *y*-axis ranges are different in Figures 2B and 6B.) Second, the network’s performance did not suffer in the Contrast condition, in which the local contrast cue was silenced (a *t*-test comparison of Thouless ratios in the Normal and Contrast conditions gave *p* = 0.23), whereas human observers’ lightness constancy was dramatically worse in the Contrast condition. Third, the network’s performance was substantially worse in the Shadows and Shading & Shadows conditions than in the Normal condition, whereas human observers performed similarly and only mildly worse, respectively, in those conditions. Fourth, differences across training instances were smaller than individual differences between human observers.

Visual inspection of the model’s output supports some of these findings, and gives insight into the model’s performance. Fig. 7 shows typical stimuli in all conditions, in a lighting configuration with strong cast shadows. The figure also shows ground-truth and network predictions for albedo and illuminance. There are clear differences in the model’s output across conditions. In the Normal and Contrast conditions, the model correctly inferred lower illuminance at the reference cube (which was in shadow) than at the match cube, and correctly assigned approximately the same albedo to the reference and match patches. In the Shadows and Shading & Shadows conditions, however, the network assigned approximately equal illuminance to the reference and match cubes, and assigned a much higher albedo to the match patch than to the reference patch. Finally, in the No-cue condition, as expected, the model inferred almost uniform illuminance everywhere, and again attributed a much higher albedo to the match patch than to the reference patch.

**Figure 7:**
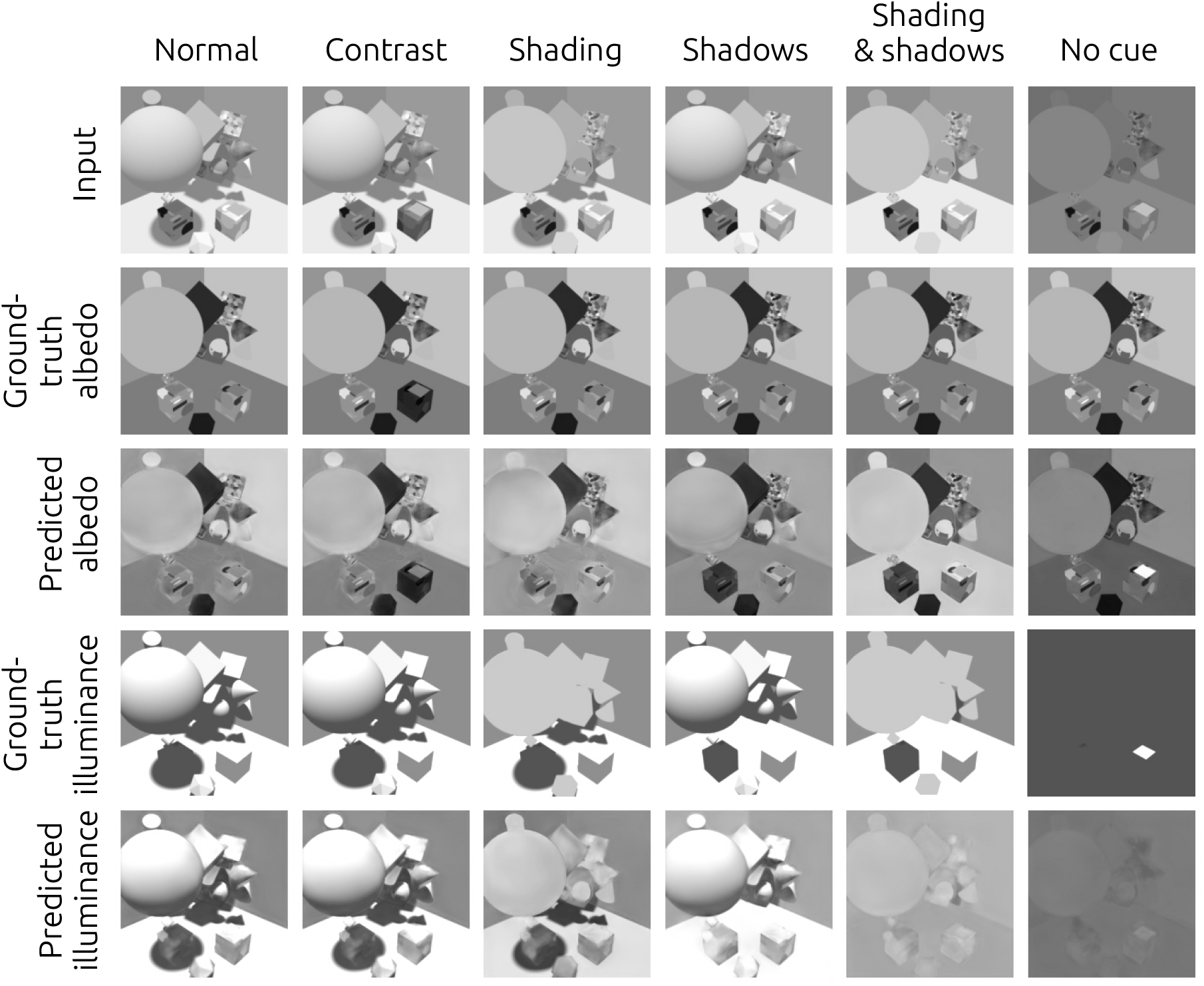
Examples of the network’s output in all six conditions. Except for the match cube in the No-cue condition, the ground-truth albedo is the same in all conditions, unlike illuminance. The model misjudged albedo in the physically inconsistent conditions. This is particularly obvious in the No-cue condition, where the model attributed the high luminance of the match patch to a high albedo instead of a high illuminance.

These results suggest that the network had some ability to model relationships over large image regions, for example using cast shadows to infer albedo at the reference and match patches. The network seemed not to rely on local contrast to estimate reflectance, but instead was able to estimate local lighting conditions, taking into account features such as shading and cast shadows. As a result, the network was susceptible to some stimulus manipulations in ways that human observers largely were not. For example, when we eliminated the shadows on the floor in the Shadows condition, the network’s estimate of local lighting conditions changed, and so did its estimates of albedo.

Overall, these results show that the network learned to use cues such as shading and shadows to disentangle the albedo and illuminance components of its input. By doing so, it achieved a much higher level of lightness constancy than human observers on the same images. It also managed to learn the non-trivial relationship between distant objects (such as the large sphere) and the shadows they cast on other objects (such as the floor and reference cube). This last feature, however, is fragile: small changes in the image can disrupt it, as when we removed the sphere’s shading in the Shading condition or the sphere’s shadow in the Shadows condition.

Most importantly, the image features that the network relies on differed from those used by human observers: while the network mostly relied on shadows, human observers’ judgements in these scenes were mostly modulated by local contrast. Thus, while the network can use naturalistic cues, its strategy for inferring albedo is quite unlike that of human observers on these images. This is consistent with the findings reported in the next section, where we demonstrate some pitfalls to be mindful of when using deep learning models of human behaviour.

## 4 Deep learning networks can exploit artifacts in rendered images

In the previous section, we saw that it is possible to train a convolutional network to disentangle the contributions of albedo and illuminance to a luminance image, relying on naturalistic features such as shading and shadows. These networks effectively learned a strategy for lightness constancy based on physically relevant cues, and one that differs from the strategy used by human observers. This was only possible due to (1) the care we took in training the models using images that, to the best of our knowledge, do not contain information other than the stipulated cues that the models could use to solve the task and (2) using test images that were novel, in the sense that these exact images were not seen by the model in training, but fell within the training distribution (i.e., they were novel, independent and identically distributed samples).

However, deep learning models, and in particular those trained via supervised learning, are prone to learn undesirable shortcuts, e.g., based on biases or artifacts in the training set (Geirhos et al., 2020). In the following section, we document that such artifacts can be present in datasets rendered with ray tracing, a popular rendering method, and how these artifacts can lead to undesirable consequences for models trained for intrinsic image decomposition.

### 4.1 Ray-tracing noise as a confounding factor

Ray tracing methods more realistic than rasterization methods because they simulate physical light transport (Shirley & Morley, 2008). They tend to be the default rendering methods used to generate photorealistic images, and in particular images used to train computer vision models. In fact, a large number of the large scale synthetic datasets available for training models for intrinsic image decomposition were rendered using ray tracing methods (Z. Li et al., 2021; W. Li et al., 2018; Zhu et al., 2022; Roberts et al., 2021). These methods’ advantage over rasterization is that they include a model of virtual light rays, *traced* back from the virtual camera to the virtual light source. However, their probabilistic nature is such that image regions with a more indirect path to the light source (e.g., regions in shadow) are less likely to be traced back, and thus tend to appear noisier. This is the case with Cycles, Blender’s ray-tracing algorithm (Blender 2.92 Documentation Team, 2021), but also many other standard ray-tracing algorithms. While there are ways to reduce the amount of noise in an image, for example by increasing the ‘sample count’ (the number of rays sampled in each pixel), areas in shadow will often nevertheless appear noisier. This correlation between illuminance and image noise is a potential confounding factor for lightness constancy, and is illustrated in Fig. 8. Thus we tested for an effect of this rendering noise in a lightness matching task, both for human observers and for the network that we examined above.

**Figure 8:**
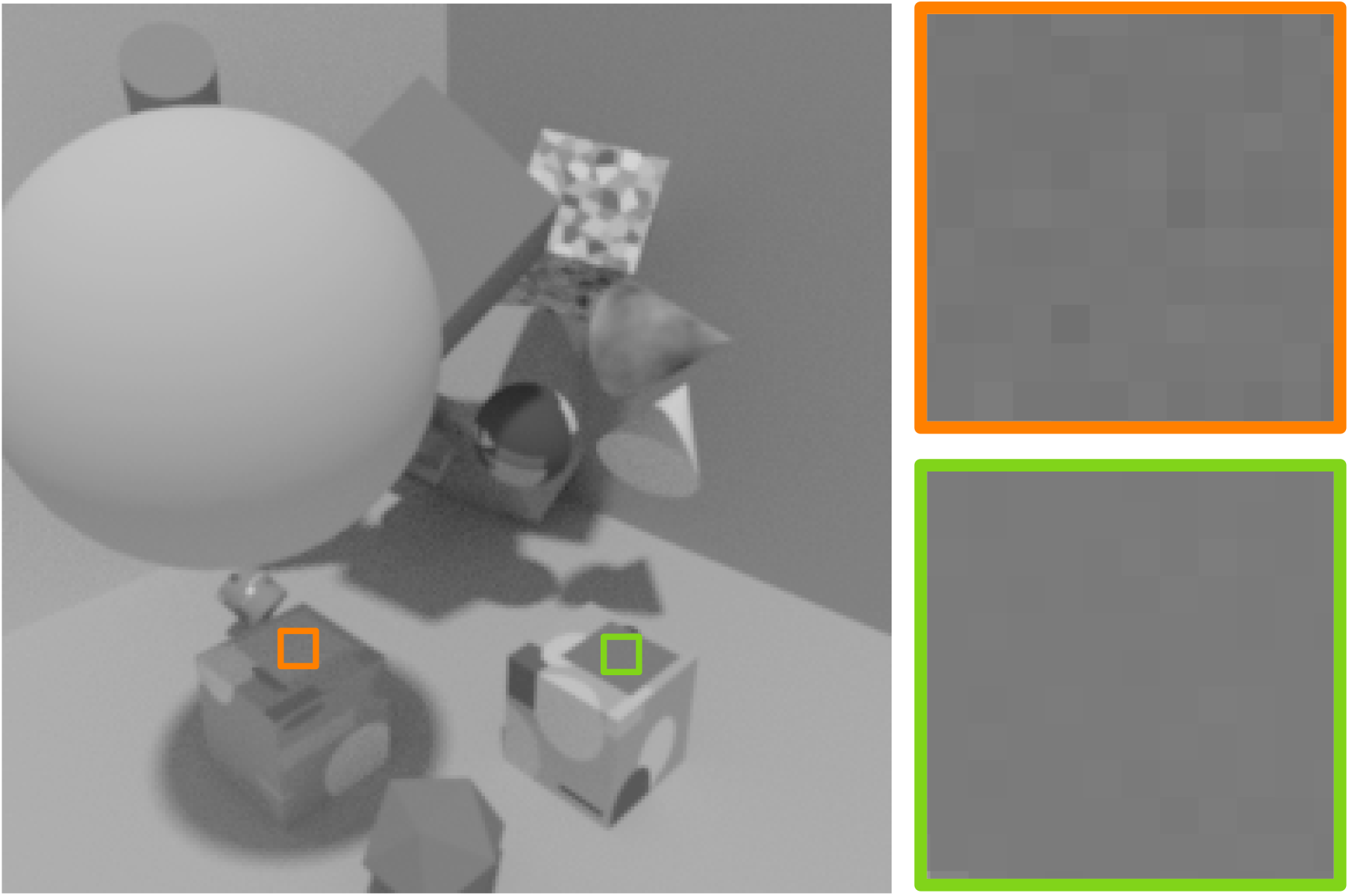
Example of image noise typically associated with ray tracing that can be a confounding factor for intrinsic image decomposition. The large squares on the right are zoomed-in views of the smaller squares in the rendered scene on the left. The two extracted patches in the image have roughly the same luminance, yet the patch in shadow (orange border) is noisier than the patch in direct light (green border). Even without a broader contextual context, this is enough local information to conclude that the patch on the left is less strongly illuminated, and thus shows an object with a higher albedo.

We rendered the same stimuli described in section 2.1, except that we used the Cycles renderer of Blender 2.92 instead of EEVEE. We used a sample count of 128, small enough that the rendering noise remains visible to the naked eye in shadowed areas. We will call this new image set the *Cycles image set*.

### 4.2 Experiment with human observers

We tested the same human participants as in the previous section, but here with the Cycles image set. As mentioned in Section 2.1, the Cycles images were interleaved with a larger number of images rendered with EEVEE, for which results are reported above. In the Cycles conditions, we tested participants only in the Normal and No-cue conditions. However, each condition included the same number of reference albedos (3), light intensities (5) and color channels converted to achromatic luminance images (3) as in the EEVEE image set. This provides a control for the main experiment: if human observers can exploit the rendering noise to perform the matching task, then there should be a significant improvement in the measured Thouless ratios in the two Cycles conditions.

That is not what we observed. There was a significant difference between the Thouless ratios measured for EEVEE and Cycles images (as shown by a repeated-measures ANOVA, *F* (1, 166) = 6.1, *p* = 0.014), but the effect was small and in the opposite of the hypothesized direction. Average Thouless ratios were 0.29 and 0.00 in the Normal and No-cue conditions, respectively, for Cycles images, compared to 0.32 and 0.04 for the EEVEE images (Fig. 9). This suggests that, if anything, the rendering noise in Cycles made the task slightly more difficult for human observers. Still, the differences are in any case very small in overall magnitude.

**Figure 9:**
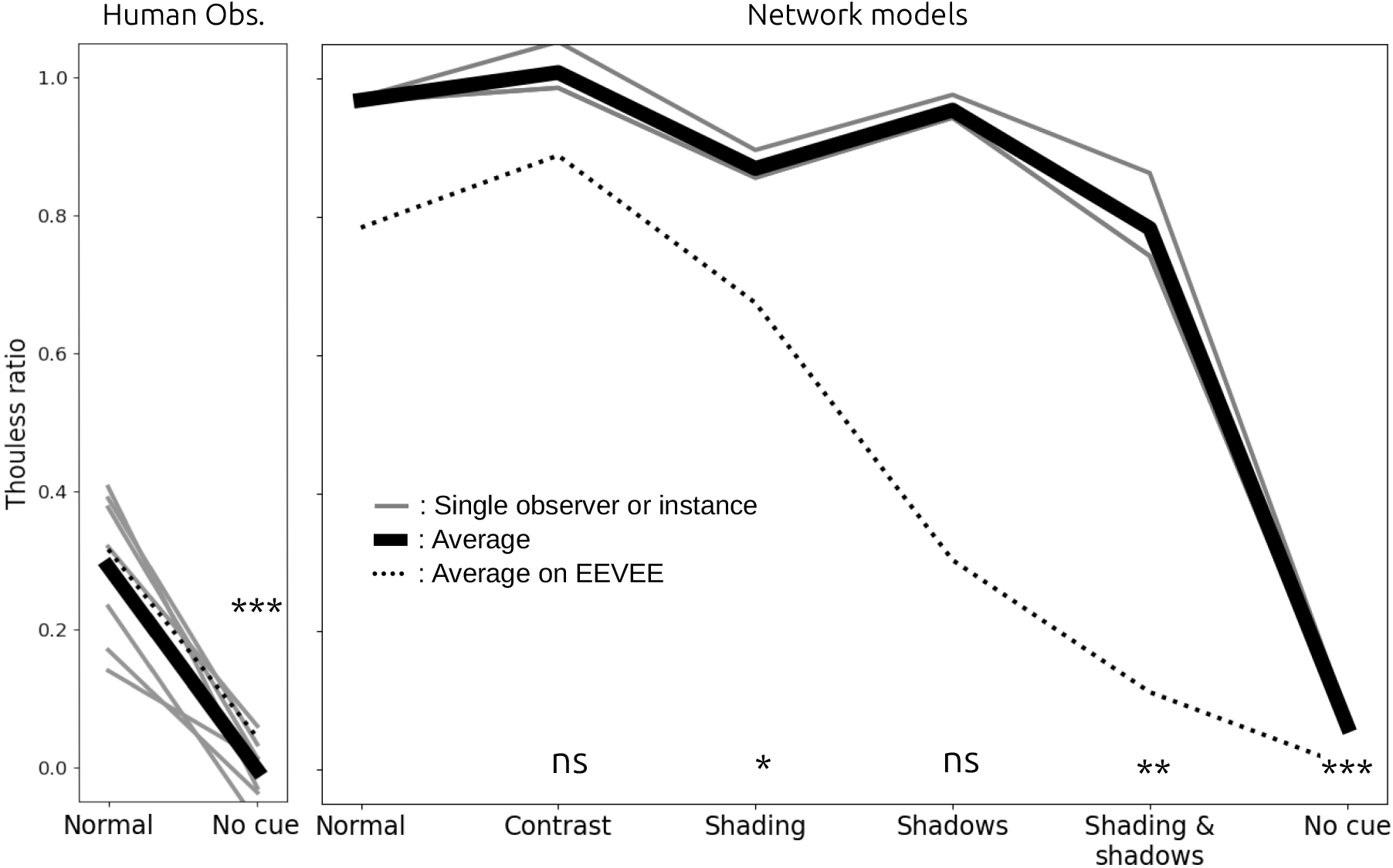
Thouless ratios, measured with the Cycles image set, for the seven human observers (left) and three training instances of the Cycles-trained model (right). Significance levels are the results of paired *t*-test comparisons to the Normal condition. The faint dashed lines are the average previously found for the EEVEE dataset, shown here for comparison purposes.

### 4.3 Deep learning models

We also tested whether DNNs trained for intrinsic image decomposition can exploit rendering noise as a cue to estimate albedo. We rendered 128K new training images and 7K new validation images with the same rendering parameters as described in Section 3.1, now using Cycles instead of EEVEE. For consistency, we used a sample count of 128. We also made three training runs of the network described above on this new image set, resulting in three training instances. We will refer to the models trained on the EEVEE images as the *EEVEE models*, and the models trained on the Cycles images as the *Cycles models*.

We tested the Cycles models on the matching task in all conditions. We found a highly significant effect of the choice of renderer on Thouless ratios (repeated-measures ANOVA, *F* (1, 106) = 19.5, *p* ≪ 0.01; Fig. 9). Thouless ratios were consistently higher for Cycles images. Additionally, although we also found a significant effect of condition on mean Thouless ratio (repeated-measures ANOVA, *p* ≪ 0.01), the Cycles models seemed to rely on different cues than the EEVEE models, as shown by the significance levels in Fig. 9 (paired *t*-test comparisons with the Normal condition). In the Normal, Contrast, and Shadows conditions, the Cycles models were essentially at perfect constancy, with average Thouless ratios of 0.97, 1.01 and 0.95, respectively, and no significant difference between them. The average Thouless ratios in the Shading and Shading & Shadows conditions were significantly lower, at 0.87 and 0.78 respectively. This suggests that the Cycles models relied on object shading, and less so on shadows, to disentangle the albedo and illuminance contribution to luminance, unlike the EEVEE models. Finally, the average Thouless ratio in the No-cue condition (0.07) was significantly greater than 0 (paired *t*-test, *p* ≪ 0.01). This last observation is particularly interesting, as in this condition there was no cue other than rendering noise that the model could use to estimate illuminance and albedo. Taken together, these findings support the hypothesis that the Cycles models learned to exploit rendering noise in order to achieve lightness constancy.

## 5 General discussion

Lightness perception is a long-standing area of research, and many studies have discovered important principles of lightness constancy (Gilchrist, 2006; Arend & Spehar, 1993; Wedge-Roberts et al., 2020; Adelson, 1993; Granzier & Gegenfurtner, 2012; Szafir et al., 2015; Murray, 2021). This research reveals lightness constancy to be a complex visual ability dependent on multiple factors, including local contrast, adaptation to the luminance range, and specular highlights, to name a few. Yet, despite these insights, few image-computable models of lightness perception exist.

Prior studies suggest that deep neural networks (DNNs) are promising candidates for such models, as they can match or even surpass human constancy levels on images of rich 3D scenes (Flachot et al., 2022; Murray et al., 2023). Our work supports this view, as our model achieved supra-human levels of constancy. However, we identified critical differences in how humans and the model solve the task. In the scenes we used, human observers relied predominantly on local contrast to judge patch colors, whereas DNNs leveraged physically relevant cues like shadows, shading, and, when present, rendering noise that was correlated with illuminance.

One could argue from these findings that unconstrained DNNs are simply too powerful to be good models of human performance in this task. Our model learned to use physically relevant features of the scenes we showed, and in order to worsen its constancy, we had to break the underlying physics by using composite scenes that silenced lighting cues from shadows and object shading. Human observers, not trained with these stimuli and possibly more limited in their capacity, relied on less computationally demanding cues like local contrast. On this view, in order to make the DNNs ‘more human’ we would have to limit their capacity, either by reducing their computational power (Kubilius et al., 2018) or by introducing additional constraints (Geirhos et al., 2018).

However, several other factors may explain the divergence between humans and the model. First, we deliberately limited the realism and complexity of the training set, in order to explore the role of generic features such as shading and occlusion. This resulted in a limited set of scene configurations (indoor), material types (Lambertian), object shapes (geometric), lighting configurations (two light sources), and shadow boundaries (sharp). Furthermore, the luminance range was limited, and in particular, specular highlights, motion, and disparity were excluded. This simplified environment made many naturalistic cues unavailable, which may have been the reason we observed low Thouless ratios for human observers. These same factors may have made shadows and shading into more reliable cues that could exploited by the network model, which was specifically trained on these scenes. One avenue for constraining networks to rely less on shadows and shading would be to make these cues more difficult to exploit, for example by increasing the complexity of the lighting, or by blurring or removing shadow boundaries. Under these conditions the network might rely more heavily on local contrast, as we found for human observers.

Second, full supervision during training is fundamentally different from how people learn to see the world (Anderson, 2015; Hoffman et al., 2015). Our visual system does not have access to the ground truth about the properties of objects and surfaces. It has been shown that fully supervised DNNs tend to develop non-human strategies, more so than weakly or unsupervised training methods (Storrs et al., 2021). It is possible that a model trained in an unsupervised manner on the same images, using for example a contrastive loss (Chopra et al., 2005), would lead to more human-like strategies. This is a promising avenue for future work.

Finally, the human visual system must solve a much broader range of complex visual tasks than just learning correlates of albedo and illuminance, such as object recognition, face recognition, and depth. Local contrast, while a valuable cue for human lightness constancy due to its invariance across changes in illuminance (Foster, 2011), is also crucial to some of these other visual functions. Indeed, early stages of the human visual system contain many contrast-sensitive cells, which are crucial for tasks like shape and object recognition (Derrington et al., 1984; Wachtler et al., 2003). Contrast-sensitive cells are also thought to be necessary for the biological system to accommodate the high dynamic range of natural images, within the smaller dynamic range of neural synapses (Shapley & Victor, 1978, 1981; Brenner et al., 2000; Gaudry & Reinagel, 2007). Contrast-sensitive cells thus form the basis of human visual processing (Hoyer & Hyvärinen, 2000). From this point of view, our observers’ sensitivity to local contrast manipulations is unsurprising. This is very different from the training environment of our network model, which was optimized solely for albedo-illuminance separation in artificial scenes where shadows and shading were highly diagnostic. Perhaps a better image-computable model of human lightness constancy would emerge from training simultaneously on a broader range of tasks, including shape and object recognition. Training such a model would require a large dataset of object-labeled naturalistic images taken under varying and known lighting conditions.

Another notable finding of the present work was the Cycles models’ reliance on ray-tracing noise which, within an image, is correlated with illuminance. In a different modeling context, Ding et al. (2019) reported the related result that if care is not taken, rendering noise can elevate the performance of trained classifiers. Given that ray-tracing is considered the most realistic rendering method (Shirley & Morley, 2008), it is the most commonly used method for generating large synthetic datasets (Z. Li et al., 2021; W. Li et al., 2018; Zhu et al., 2022; Roberts et al., 2021). As a result, many models trained for inverse rendering and intrinsic image decomposition are trained on ray-traced datasets (Z. Li & Snavely, 2018a; Z. Li et al., 2020; Wang et al., 2021; Zhu et al., 2022) and may be susceptible to similar artifacts. This could limit their performance when they are applied to natural images, in which such artificial noise is not present. This issue is particularly relevant for models of human lightness and color constancy. To mitigate it, we suggest avoiding ray-tracing algorithms in favor of methods like rasterization, or potentially mixing ray-traced and rasterized images in training. Alternatively, recent years have seen the development of powerful de-noising methods (Bako et al., 2017; Chopra et al., 2005) and these methods might silence rendering noise as a cue for disentangling albedo and illuminance. Regardless, in experiments that evaluate network performance, particularly when trained and evaluated on rendered scenes, we recommend including control conditions where good performance should be impossible, like our No-cue condition, as a test for the presence of unwanted cues.

Overall, our results show deep learning networks to be a promising avenue for developing models of human lightness constancy. There are few image-computable models of human lightness perception, and the ones that have been developed fail often in complex scenes (Figure 4). Here we find the opposite problem: DNNs can succeed even where human observers fail. However, this does not discredit the potential of DNNs to model human lightness constancy, and human vision in general. DNNs remain a powerful class of image computable models, and our findings show that when images are carefully selected to exclude artifacts, DNNs can exploit relevant, naturalistic cues like shadows and shading. There are several factors that may explain the divergence we found between the network model and human observers, including network capacity, the realism of the training set, the kind of training, and the need to simultaneously learn several visual tasks. Exploring these factors is a promising approach for future work.

## Acknowledgements

This work was funded by the following sources. (a) A postdoctoral fellowship to AF from the Canada First Research Excellence Fund, through the Vision: Science to Applications program at York University. (b) NSERC Canada Graduate Scholarship - Doctoral to JP. (c) European Union funding to TSAW (ERC, SEGMENT, 101086774). Views and opinions expressed are however those of the author(s) only and do not necessarily reflect those of the European Union or the European Research Council. Neither the European Union nor the granting authority can be held responsible for them. This research was cofunded by Research Cluster “The Adaptive Mind”, funded by the Excellence Program of the Hessian Ministry of Higher Education, Science, Research and the Arts. (d) An NSERC Discovery Grant to RFM. (e) Computing resources were provided by SHARCNET (www.sharcnet.ca) and the Digital Research Alliance of Canada (alliancecan.ca).

## Footnotes

1 https://github.com/AlbanFlachot/IID4constancy/tree/main

